# Why are there so many independent origins of artemisinin resistance in malaria parasites?

**DOI:** 10.1101/056291

**Authors:** Timothy J.C. Anderson, Shalini Nair, Marina McDew-White, Ian H. Cheeseman, Standwell Nkhoma, Fatma Bilgic, Rose McGready, Elizabeth Ashley, Aung Pyae Phyo, Nicholas J. White, François Nosten

**Affiliations:** Department of Genetics, Texas Biomedical Research Institute, San Antonio, Texas, United States of America; Shoklo Malaria Research Unit, Mahidol-Oxford Tropical Medicine Research Unit, Faculty of Tropical Medicine, Mahidol University, Mae Sot, Thailand; Mahidol-Oxford Tropical Medicine Research Unit, Faculty of Tropical Medicine, Mahidol University, Bangkok, Thailand; Centre for Tropical Medicine, Nuffield Department of Medicine, Churchill Hospital, University of Oxford, United Kingdom

## Abstract

**Summary:** Multiple alleles at the *kelch13* locus conferring artemisinin resistance (ART-R) are currently spreading through malaria parasite populations in Southeast Asia, providing a unique opportunity to directly observe an ongoing soft selective sweep, to investigate why resistance alleles have evolved multiple times and to determine fundamental population genetic parameters for Plasmodium. We sequenced the *kelch13* gene (n=1,876), genotyped 75 flanking SNPs, and measured clearance rate (n=3,552) in parasite infections from Western Thailand (2001-2014). We describe 32 independent coding mutations: these included common mutations outside the *kelch13* propeller region associated with significant reductions in clearance rate. Mutations were first observed in 2003 and rose to 90% by 2014, consistent with a selection coefficient of ~0.079. There was no change in diversity in flanking markers, but resistance allele diversity rose until 2012 and then dropped as one allele (C580Y) spread to high frequency. The rapid spread of C580Y suggests that the genomic signature may be considerably harder in the near future, and that retrospective studies may underestimate the complexity of selective sweeps. The frequency with which adaptive alleles arise is determined by the rate of mutation to generate beneficial alleles and the population size. Two factors drive this soft sweep: (1) multiple amino-acid mutations in *kelch13* can confer resistance providing a large mutational target – we estimate the target size is between 87 and 163bp. (2) The population mutation parameter (*Θ*=2*N_e_μ*) can be estimated from the frequency distribution of resistant alleles and is ~ 5.69, suggesting that short term effective population size is between 88 thousand and 1.2 million. This is 52 to 705-fold greater than *N_e_* estimates based on fluctuation in allele frequencies, suggesting that we have previously underestimated the capacity for adaptive evolution in Plasmodium. Our central conclusions are that retrospective studies may underestimate the complexity of selective events, ART-R evolution is not limited by availability of mutations, and the *N_e_* relevant for adaptation for malaria is considerably higher than previously estimated.

**Significance Statement:** Previous work has identified surprisingly few origins of resistance to antimalarial drugs such as chloroquine and pyrimethamine. This has lead to optimism about prospects for minimizing resistance evolution through combination therapy. We studied a longitudinal collection of malaria parasites from the Thai-Myanmar border (2001–14) to examine an ongoing selective event in which ≥32 independent alleles associated with ART-R evolved. Three factors appear to explain the large number of origins observed: the large number of amino acid changes that result in resistance (i.e. large mutational “target size”), the large estimated effective population size (*N_e_*), and the fact that we were able to document this selective event in real time, rather than retrospectively.

## INTRODUCTION

The number of times drug resistance alleles arise is a central question for disease intervention because it determines the useful life of drugs, the ease with which resistance arises, and the utility of diagnostic markers for tracking resistance. It is also a question of central interest for evolutionary biologists interested in adaptation, where the relative role of hard selective events, in which flanking genetic variation is purged from around the selected site, and soft selective events in which flanking genetic variation is maintained, is being actively debated (1, 2). The frequency with which beneficial alleles establish is expected to be determined by the rate (*μ*) at which mutations conferring resistance are introduced into the population (by mutation or migration) and the effective size (*N_e_*) of the population during the time when selection is acting: these parameters are combined in the population mutation parameter (*Θ* = 2*N_e_μ*). Elegant theoretical work (3, 4) has led to the prediction that when *Θ*<*0.01*, mutations will be limiting and most selective events will involve single alleles generating hard selective sweeps, when *Θ* is between *0.01* and *1* both single and multiple origins of adaptive alleles are possible, and when Θ > 1, multiple origins of adaptive alleles are expected.

Most studies of selection in nature are retrospective, so the underlying dynamics of these events are incompletely understood. Typically, genetic data from present day populations is used to infer selective events that have occurred in the past. The surge of recent next generation sequencing studies has generated large genomic datasets from multiple organisms and haplotype tests for detecting signatures of past selection have been widely used to catalogue selected genome regions. However, these methods can only detect the “fossil record” of selection in the genome. While valuable, this genomic fossil record may provide a simplified description of adaptive evolution, which is biased because only surviving allelic lineages are examined. In contrast, molecular description of ongoing adaptive evolution in nature provides a direct record of the dynamics of selective events, including documentation of allelic lineages that fail to survive. Molecular dissection of rapid evolution of microbial or insect populations in the face of ongoing selection with drugs of pesticides are particularly valuable in this respect (5–8).

Malaria parasites provide a powerful system for studying adaptation in nature, because parasite populations have been exposed to a succession of antimalarial compounds over the past 75 years, and selection is strong, with coefficients of 0.1-0.4 recorded (9). Parasite genomes are compact (23Mb), blood stage parasites are haploid, and there is an obligate recombination in the mosquito stage, making this a useful system for understanding adaptation in other recombining eukaryotes. Retrospective molecular studies of resistance to chloroquine and pyrimethamine, in which variation at markers flanking resistance mutations were examined, have revealed remarkably few independent origins of resistance alleles (10–12). Selection appears to have driven textbook “hard” selective sweeps, in which single resistance haplotypes have rapidly spread through malaria parasite populations purging genetic variation in the vicinity of the resistance locus and spreading across global parasite populations. The rarity of independent resistance alleles is surprising given that each infected human contains up to ~10^11^ parasites, so we would expect that each infected person will contain 10-100 parasites with mutations at each position in the genome assuming standard eukaryotic SNP mutation rates. However, the rarity of mutations is consistent with measures of short-term *N_e_*, determined from temporal fluctuation in allele frequencies: these genetic-drift based *N_e_* estimates range from under one hundred (13) to several thousand (14). However, these studies of chloroquine and pyrimethamine resistance were retrospective (15) because they examined parasite population samples collected ~25 years after the origins of resistant parasites in the 1950-70s. Hence we were unable to precisely document these selective events as they were occurring.

The recent discovery of a major ART-R gene, *kelch13*, in SE Asia, in which resistance alleles clearly have multiple independent origins (16–18), supports a radically different model of resistance evolution, and provides a unique opportunity to examine an ongoing selective event in a recombining eukaryote and to estimate key population parameters for *Plasmodium*. Transfection experiments unambiguously demonstrate that single amino acid edits in *kelch13* increase survival of cultured parasites (19), showing that this gene alone is sufficient to cause ART-R. ART-R was first confirmed in SE Asia in 2009 (17). ART-R parasites show 100-1,000 fold slower clearance than sensitive parasites over a single 48 hour parasite life-cycle, and this results in increased treatment failure rates (20–22). Here, we describe the spread of resistance alleles in a longitudinal collection of parasites from the Thailand-Myanmar border between 2001 and 2014. Thirty-two independent mutations have spread through this parasite population generating an extraordinarily soft selective event. However, haplotypes associated with one particular mutation now dominate this parasite population, so the dynamics of this sweep are soft, but the signature in several years time is expected to be that of a much harder sweep. We use these data to estimate critical parameters such as the population mutation rate (3, 4) and the short term effective population size (*N_e_*) for malaria and more generally to ask why there should be so many independent origins of ART-R.

## MATERIALS AND METHODS

### Study site and patients

We collected *P. falciparum* infected blood samples from patients visiting malaria clinics run by the Shoklo Malaria Research Unit (SMRU) which span a 150 km region of the North-Western border of Thailand between 2001 and 2014. The blood samples collected were finger prick blood spots (~30μl whole blood dried on Whatmann 3M filter paper) or venous blood samples depleted of white blood cells (23). The four clinics sampled, which span ~100km along the western border of Thailand, were Maela (refugee camp) and Wang Pha village to the north of Maesot, and Mae Kon Khen and Mawker Thai villages to the south of Maesot. The majority of patients came from adjacent Myanmar. Falciparum malaria was diagnosed by microscopy of thick and thin peripheral blood smears stained with Giemsa. Parasite counts were read per 1,000 red cells (thin film) or 500 white cells (thick film). The parasite samples included in this study came from three different groups of patients: (a) Uncomplicated hyperparasitaemia patients with ≥4% of red cells infected. These patients were treated with artesunate (ART) alone followed by combination therapy as described previously (24). Following treatment, we monitored parasite clearance by measuring parasite density in the blood at 6 hourly intervals until the patient became slide negative. These data were used to measure parasite clearance rates (see below). (b) We collected pre-treatment blood samples from patients with uncomplicated malaria (<4% of red cells infected) included in drug efficacy studies between 2003-2013 and treated with mefloquine+artesunate (22). Parasite clearance rates were not measured in these patients. (c) Numbers of *P. falciparum* cases are dropping rapidly in this region (14, 25). In 2014, we therefore supplemented the numbers of samples for molecular analysis by collecting finger prick blood spots from malaria patients routinely admitted for treatment to these four clinics. Ethical approval for this work was given by the Oxford Tropical Research Ethics Committee (OXTREC 562-15) and the Faculty of Tropical Medicine, Mahidol University (MUTM 2015-019-01).

### Measurement of parasite clearance half-life

Slow parasite clearance from the bloodstream of infected patients is a hallmark of ART-R (26) and was determined in all hyperparasitaemia patients. We measured the rate of parasite clearance by plotting log parasite density against time using a standardized fitting method which separates the variable initial lag-phase from the subsequent log-linear decline (27) using the online parasite clearance calculator (http://www.wwarn.org/tools-resources/toolkit/analyse/parasite-clearance-estimator-pce). We measured the slope of the linear phase and determined the parasite clearance half life (*T_1/2_P*), defined as the time required for parasitaemia to fall by half during the log-linear decline. We excluded parasite clearance curves showing a poor fit (r^2^ < 0.8) to the log-linear model.

### Genotyping of parasites

We extracted DNA admission blood spots using a two step protocol to maximize DNA yield. Blood was eluted from the six three mm punches using the Gensolve kit (GenVault Corporation) and DNA extracted using 96-well QIAamp 96 DNA Blood Kits (Qiagen). Whole blood samples were extracted using the Gentra PureGene kit (QIAGEN) and DNA concentration was quantified using a Qubit fluorometer.

We genotyped parasite infections from hyperparasitemic patients using 96-SNPs distributed across the *P. falciparum* genome using the Illumina Veracode platform. We considered infections to contain multiple clones if >5 SNPs showed heterozygous base calls (14). We used single clone infections for subsequent molecular analyses, because haplotypes can be unambiguously constructed.

To further assess haplotype structure around the *kelch13* locus, we genotyped an additional 163 SNPs using Goldengate from a subset of samples (genetically unique single clone infections from the hyperparasitemia study, and all samples from the treatment efficacy study). The 163 SNPs comprised included 75 SNPs on chr 13 as well as 88 SNPs on other chromosomes. The SNPs genotyped were located in genome regions showing association with ART-R in previous association (28) and pooled sequencing studies (29), so are not randomly ascertained.

### Sequencing of *kelch13* locus

We amplified three PCR fragments from the *kelch13* locus and sequenced products in both directions on an ABI 3730 capillary sequencer: fragment 1 (1,725,980-1,726,520bp, pos 419-570): F- ATCTAGGGGTATTCAAAGG, R- CCAAAAG ATTT AAGT G AAAG; fragment 2 (1,726,400-1,726,940bp, pos 545-707): F- CTGCCATTCATTTGTATCT, R- GGATATGATGGCTCTTCTA), fragment 3 in the 5′ region (1,725,380-1,725,680bp, pos 211-302): F- T GAAAAT ATGGTAGGT GATT and R- ATCGTTTCCTATGTTCTTCT. We treated PCR products with Exo-SAP-IT (GE Healthcare), and sequenced them directly in both directions using the BigDye Terminator v3.1 cycle sequencing kit (Applied Biosystems, Inc., Foster City, CA). We cleaned BigDye products using the BigDye XTerminator Purification kit (Applied Biosystems, Inc.) and ran them on an ABI 3730 capillary sequencer. We aligned and analyzed data using SeqScape version 2.7. We excluded parasite samples with incomplete sequence data for one or more fragments from the analysis.

### Analysis

All analyses were conducted using R version 3.2.3.

<u>Selection coefficients</u> – We measured selection coefficients for *kelch13* mutations by plotting log(R/S) over time and measuring the slope. Time was measured in *Plasmodium* generations (the time taken for parasites to traverse both mosquito and human stages of the lifecycle). The length of a *Plasmodium* generation varies and is dependent on multiple factors such as gametocyte production and vector abundance. We used generation times of 1.5, 2 and 3 months.

<u>Modeling the selective sweep</u> – Whether selective sweeps are soft or hard is critically dependent on the rate at which beneficial mutations are introduced into the population, which is determined by the population size over the time period in which selection is acting, and the rate at which beneficial mutations are generated. We used published theory (3, 4) to examine the fit of our data to predictions for soft selective sweeps. The critical parameter determining whether soft selective sweeps will occur is the population mutation parameter (*Θ*=2*N_e_μt*), where *N_e_* is the effective population size during the time period in which adaptation occurs (here between 1995 when ART treatment was first introduced and the present time where resistance alleles are close to fixation), *μ* is the per nucleotide mutation rate per *Plasmodium* generation (the complete lifecycle from including mosquito and human development), and *t* (the mutational “target” size) defined as the number of nucleotides at which mutations can generate a resistance phenotype.

*Θ* can be determined directly from the numbers of independent origins of resistance mutations in *kelch13* in our population sample using eqn. 12 in Pennings and Hermisson (3):

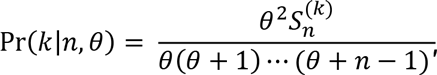
where 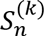 is Stirling’s number of the first kind, *n* is the sample size and *k* is the number of haplotypes observed. We determined 95% confidence intervals from the Pr(*k*|*n,θ*) probability distribution across a range of *Θ values*. Since *Θ*=2*N_e_μt*, we estimated *N_e_* using measures of mutation rate (*μ*) and target size (*t*): *N_e_* =*Θ*/2*μt*.

The mutation rate (*μ*) has been measured in two mutation accumulation experiments (30, 31). These estimates range from 1.7−3.2 × 10^−9^ (31) to 3.82−4.28 x×10^−10^ per base/48hr asexual cycle (30). The estimates are mutation rates for a single 48 hour asexual parasite cycle. We use the mean mutation rate (3.39 ×10^−9^, range: 1.7−4.28 ×10^−9^) and converted per cycle rates to per generation mutation rates by multiplying by *g/2*, where *g* is the length in days of a malaria parasite generation (the duration of the complete parasite lifecycle, from mosquito to mosquito), and 2 days is the length of a single asexual cycle. We use a two month generation time per year so g=~60 days. We have assumed that the rate of cell division is the same in mosquito and human sections of the parasite lifecycle.

We estimated the mutational target size (*t*) – the number of bases at which mutations can generate ART-R – by counting the numbers of different mutations associated with ART-R in this study and in other recent papers describing sequence variation in *kelch13*. We then used rarefaction approaches (32) to estimate the number of possible resistance alleles, from the rate at which new alleles are discovered.

<u>Genotype-phenotype associations</u> – We compared clearance half life (T*_1/2_*P) of parasites bearing *kelch13* mutations compared with wild-type alleles using t-tests. We used uncorrected tests to avoid needless rejection of the null hypothesis (i.e. no difference in T_1/2_P) due to low sample size (8 mutations are found in ≤3 samples), and because the vast majority of mutations are expected to differ in phenotype from WT. In addition, we examined pair wise differences in *T_1/2_P* between different *kelch13* alleles to determine allele specific variation in clearance: in this case Bonferroni correction was used.

<u>SNP variation on chromosome 13</u> – We measured extended haplotype homozygosity (EHH) using *rehh* (33) to examine hitchhiking associated with different *kelch13* alleles. We computed SNP-π for each bi-allelic SNP: this is defined as the average proportion of pairwise differences at assayed SNP loci within a defined population (34). Average SNP-π across SNPs was then computed for SNPs flanking *kelch13* to investigate changes in genetic variation over time. We measured expected heterozygosity (*H_e_*) at the *kelch13* locus by treating *kelch13* as a single locus with multiple alleles. To evaluate evidence for independent origins of particular ART-R mutations, we used *poppr* (35) with default settings to construct minimum spanning networks comparing the relationships between different SNP haplotypes for the chr. 13 region surrounding *kelch13*. This analysis used chr. 13 data with ≤5 missing SNPs or ambiguous base calls.

## RESULTS

### Dataset

We collected parasite genotypes, *kelch13* sequence and/or parasite clearance data from 4,867 patients visiting four clinics along the Thailand-Myanmar border between 2001-2014. These patients included 3,878 hyperparasitaemia patients, 917 patients enrolled in an efficacy study of ART-mefoquine, and 72 infections from uncomplicated malaria patients visiting SMRU clinics. Fig. S1 shows how these patient samples were used for different analyses. After removing data whose clearance rate curve fitted poorly to a linear model (r^2^≥0.8) 3,552 remained. We excluded multiple clone infections by genotyping a panel of 93 variable SNPs (24) in the hyperparasitemia samples. We determined sequence at the *kelch13* locus in 1,876 infections. In addition, to examine the impact of *kelch13* on flanking variation we genotyped an additional 163 SNPs (75 on chr. 13, and 88 elsewhere in the genome) in a subset of samples.

### Change in parasite clearance rate

We previously described the change in clearance rate from 2001-2010 (24). Mean *T_1/2_P* accelerates during 2011-14, reaching over 6.0hrs in 2014 (Fig. 1A). The dynamics of CR increase differ between locations (Fig. S2).

### Mutations in the *kelch13* locus

We observed 32 independent non-synonymous mutations in the *kelch13* locus. The 32 mutations include 29 in the propeller region of the gene (1725980-1726940bp, amino acid positions 419-707), as well as 3 outside the propeller (E252Q, D281V, R239Q). We observed multiple mutations in several codons. For example P667→Q, R or T (CCA→C<u>A</u>A, C<u>G</u>A, or <u>A</u>CA), while at P527 consecutive mutations have generated H or L (CCT→C<u>A</u>T→C<u>T</u>T). Mutations were first observed in 2003: the percentage of parasites carrying mutations at *kelch13* rose to 90% in 2014 (Fig. 1B). The rise in *kelch13* allele frequency closely tracks change in parasite *T_1/2_P* over time in the 4 clinics examined (Fig. 1A and B, S2 and S3) and we see parallel changes in both hyperparasitemia and efficacy study data sets (Fig. S4). The most abundant allele before 2008 was E252Q (Fig. 1B and D). C580Y allele was first observed in 2006, reached 4% frequency in 2010, and by 2014 reached 65% (Fig. 1B and D) (72% of resistant alleles) and was at fixation or close to fixation in all clinics except for MLA (Fig. S3). Only two other alleles reached over 10% allele frequency (E252Q in 2007 and 2009-12, and R561H in 2012) during the 14 year sampling period.

**Fig. 1.**
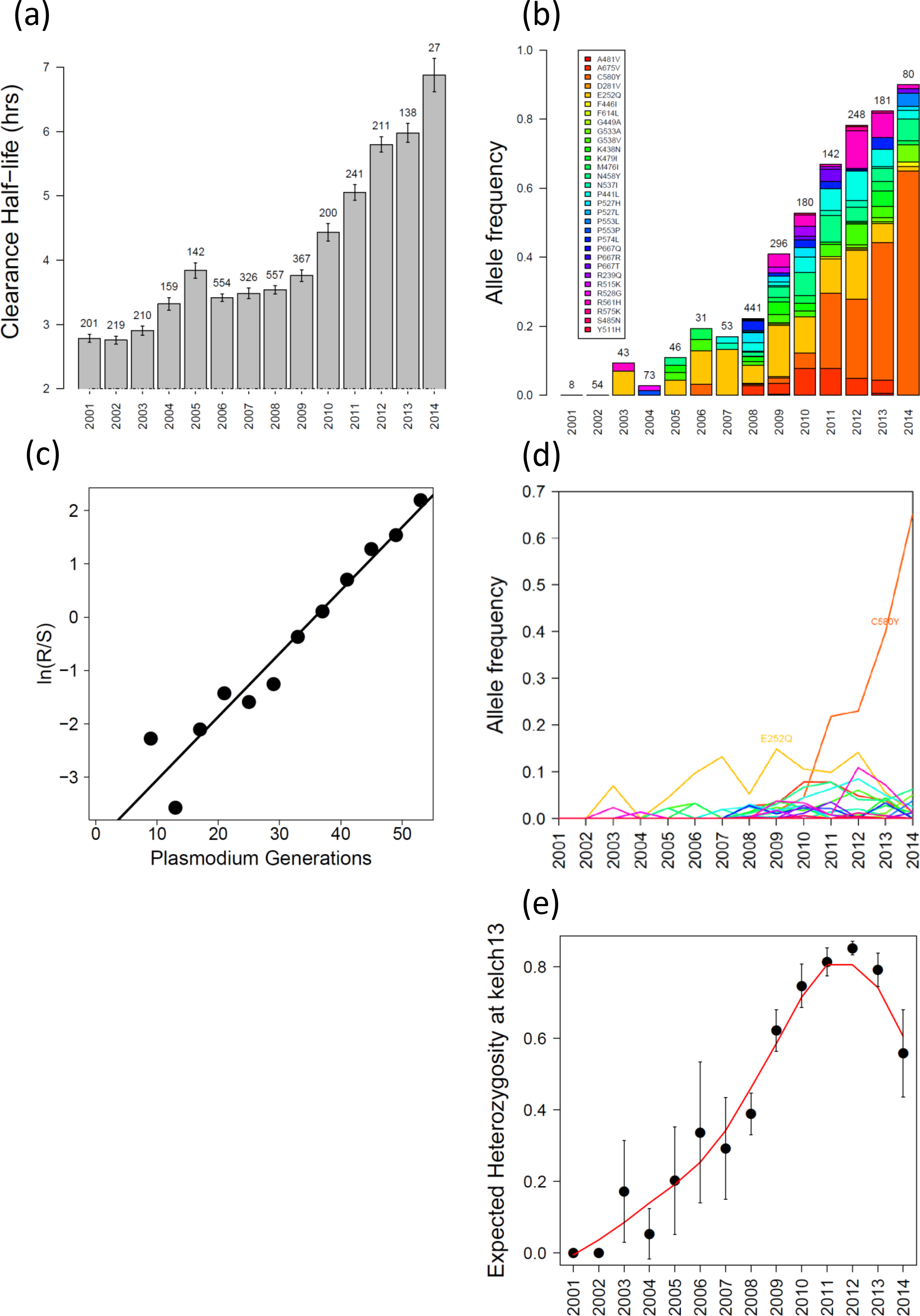
Evolution of ART-R on the Thailand-Myanmar border. A. Change in clearance rate (*T_1/2_P*) between 2001 and 2014. B. Frequency of alleles carrying different *kelch13* mutations on the Thailand-Myanmar border. Infections carrying a single predominant infections (assessed using SNP genotyping) were studied. Sample sizes are shown above the bars, and colors indicate the frequency of different alleles (see key). The graph includes samples from both hyperparasitaemia (single genotype infections) and drug efficacy studies (all infections): these datasets are plotted independently in Fig. S4. C. Change in frequency of *kelch13* alleles. Lines show the trajectory of different *kelch13* alleles over time. E252Q predominates initially, while C580Y spreads to high frequency after 2011. D. Selection coefficient estimation for resistance alleles on the Thailand-Myanmar border. E. Expected heterozygosity (*H_e_*) of *kelch13* alleles over time. This drops rapidly from a peak in 2012.

We determined selection coefficients driving *kelch13* mutations (Fig. 1C) by assuming 4, 6 or 8 parasite generations per year. Selection coefficients were 0.12, 0.079, and 0.059 respectively. The analysis treats all alleles as being equivalent. The very rapid rise of C580Y indicates much higher allele-specific selection coefficients: 0.24, 0.16 and 0.12 assuming 4, 6 or 8 parasite generations per year.

We measured expected heterozygosity (*H_e_*) at *kelch13* (the probability of drawing two different alleles). This rises from 0 in 2001 to 0.8 in 2011 and then drops to 0.5 as C580Y spreads in the population (Fig. 1E).

### Genotype-phenotype associations

We compared *T_1/2_P* in parasite infections carrying different *kelch13* alleles (n=1,044, Fig. 2). Parasites with WT *kelch13* alleles show a low mean CR, and were cleared from the bloodstream significantly faster than parasites with mutations in *kelch13* (t = −34.99, 3.77 × 10^−165^). The distribution of CR in WT parasites was broad (1.065-10.21h, median=2.86h): however just 26/606 (4.2%) WT parasites had *T_1/2_P* ≥ 5h. We observed significant variation in *T_1/2_P* associated with different *kelch13* alleles (Fig. 2A). Two non-synonymous SNPs (K438N and F614L) showed no association with elevated clearance rate. After removing these and WT alleles, we observed significant heterogeneity in impact of mutations on *T_1/2_P* (F=1.29, df=22, p < 6.21 × 10^−31^). All three mutations outside the propeller region (R239Q, E252Q, and D281V) were associated with moderate *T_1/2_P* (4.53, 4.57, and 5.41hr respectively): parasites bearing these mutations cleared significantly slower than WT (t = −14.44, 5.58 × 10^−30^), and significantly faster than parasites bearing C580Y (t = 11.77, 4.18 × 10^−25^). C580Y (*T_1/2_P* = 6.35) parasites cleared faster than several other *kelch13* common ART-R alleles (R561H (*T_1/2_P* = 7.00) and N458Y (*T_1/2_P* = 7.12)), although these differences were not significant after correction for multiple testing (Fig. 2B).

**Fig. 2.**
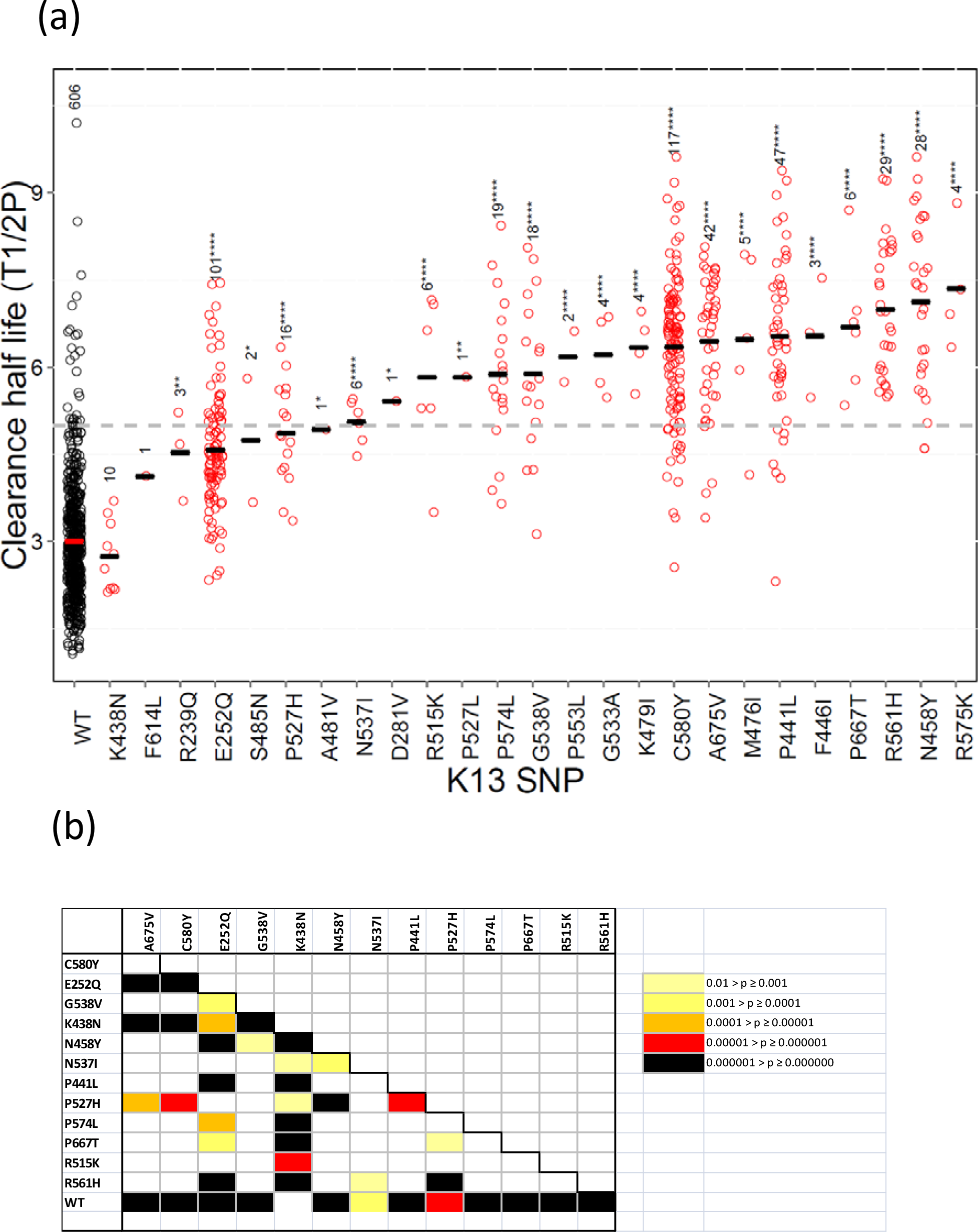
Association between *kelch13* alleles and parasite clearance phenotype. A. Each circle represents *T1/2P* from a single patient. Sample sizes are shown above the highest value for each mutation, and median values are shown. The horizontal dotted line at T _1/2_P=5 provides a commonly used cut-off for resistant infections used for categorizing clinical infections. Asterisks mark significant differences from WT alleles (* p<0.05;** p<0.01; *** p<0.001; **** p<0.0001) using t-tests. Uncorrected tests were used to avoid needless rejection of the null hypothesis due to low samples size (8 mutations are found in ≤3 samples), and because the vast majority of mutations are expected to differ in phenotype from WT. All comparisons are significant at α = 0.05 threshold except K438N and F614L. B. Significant variation between *kelch13* alleles in phenotypic impact on *T_1/2_P*. Bonferroni corrected pairwise comparisons of *T_1/2_P* between parasites bearing different *kelch13* mutations. Note that parasites bearing E252Q mutations (outside the propeller region of the gene) show significantly slower clearance than WT parasites, but significantly more rapid clearance than parasites with mutations in the *kelch13* propeller, such as C580Y or R561H.

Whether additional loci in the genome impact clearance rates is an open question. We examined whether clonally indistinguishable groups of parasites isolated from different patients, but bearing the same mutation showed significant heterogeneity in *T_1/2_P* for common genotypes including WT, E252Q, and C580Y. Among the 612 WT parasites there were 88 groups of unique WT genotypes infecting 2-13 patients, but we observed no evidence for heterogeneity in *T_1/2_P* (F=0.0373, df=, ns). Similarly for C580Y (n=99, 15 groups infecting 2-6 patients, F=0.031, df=, ns) and E252Q (n=00, 18 groups infecting 2-10 patients, F=.1275, df=, ns) we found no evidence for genetic background effects influencing T*_1/2_*P (Table S1).

### Linkage disequilibrium and hitchhiking around *kelch13*

Soft sweeps are expected to have much less impact on flanking diversity than hard sweeps, because multiple background haplotypes hitchhike with selected mutations. We measured SNP diversity (*SNP*-π) surrounding WT and alleles associated with elevated clearance rate for 2008-13. We observed no difference in diversity between parasites carrying WT and *kelch13* ART-R alleles on any year examined (Fig. 3A). However, when we examined *SNP*-π for SNPs surrounding individual *kelch13* SNPs alleles we observed significant reduction in diversity relative to WT parasites for all common SNP alleles except M461I and E252Q (Fig. 3B). This was also reflected in increased linkage disequilibrium surrounding *kelch13* ART-R alleles (Fig. 3C–D, Fig. S5). We inspected the length of haplotypes surrounding individual *kelch13* alleles from the decay in expected haplotype homozygosity (EHH), defining elevated EHH as above 0.2. All common alleles examined showed elevated EHH (432-2,130kb) relative to WT alleles (182kb) (Fig. 3D). Two particular alleles showed extremely long tracts of high EHH (G538V: 2446kb; P441L: 2130kb). E252Q has the shortest region of elevated EHH (432kb), but there was no relationship between the impact of alleles on clearance rate, and the length of LD observed. Furthermore, C580Y, which is increasing in frequency rapidly in this and other SE Asian populations, showed moderate levels of LD.

**Fig. 3.**
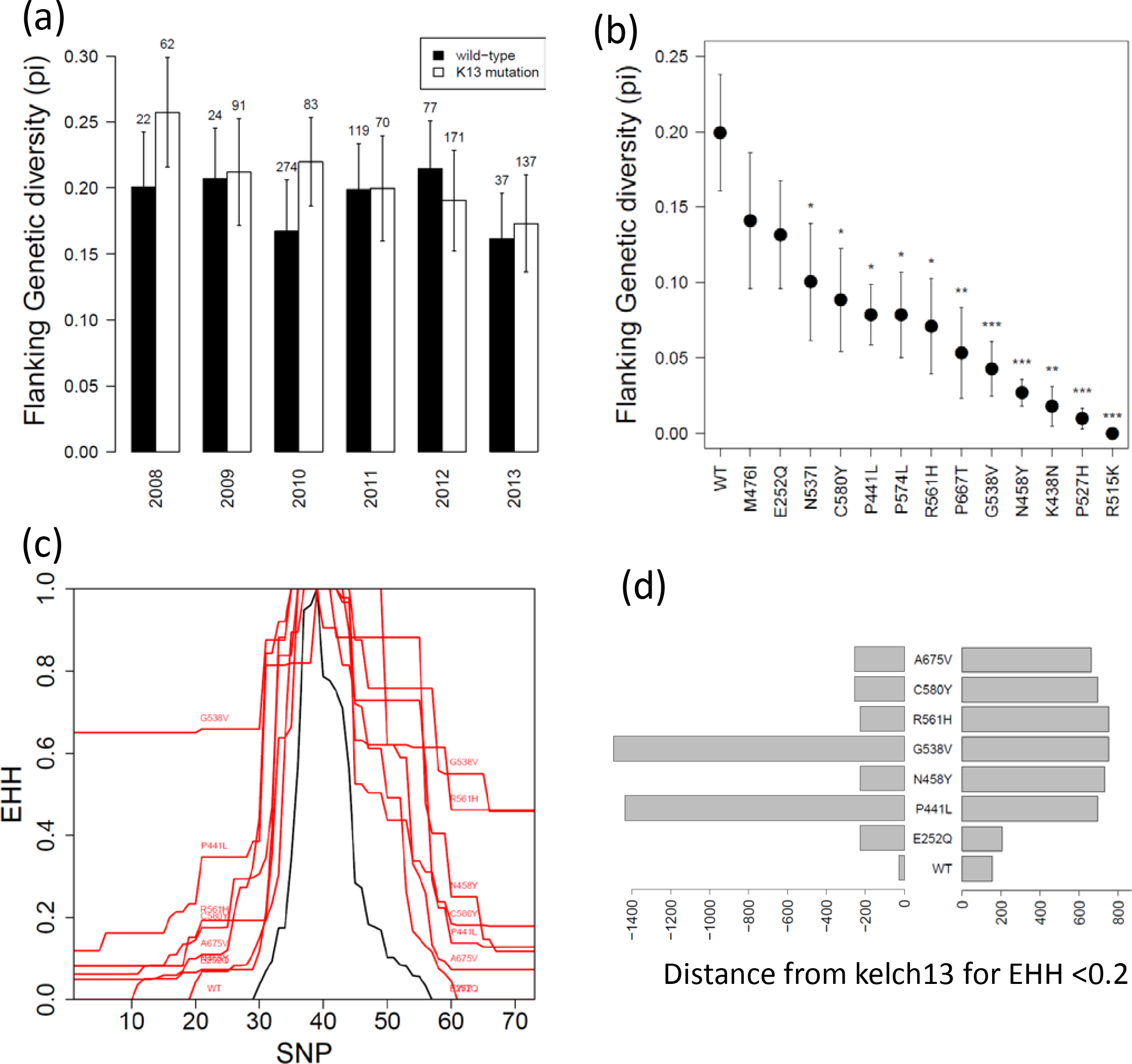
Genetic diversity and linkage disequilibrium around resistance alleles. A. Mean diversity (*SNP*-π) for genotyped SNPs flanking *kelch13* on WT (black bars) and *kelch13* alleles carrying mutations (white bars). We observed not difference in diversity between these two groups. B. Mean diversity (*SNP*-π) surrounding individual *kelch13* alleles. All alleles except M476I and E252Q show significantly reduced variation relative to WT alleles (Bonferroni correct p-values: * p<0.05, ** p<0.01, *** p<0.001).C. Expected haplotype homozygosity (EHH) plotted across chr. 13. Black line shows EHH surrounding WT alleles, while the red lines show EHH surrounding common *kelch13* alleles. The markers are unevenly spaced on chr. 13 (fig. S5), but are ordered on the x-axis: D. Length of extended haplotypes surrounding *kelch13* alleles. Bars show the distance at which EHH decays below 0.2 on either side of *kelch13* on chr. 13.

### Multiple origins of individual resistance mutations

*Kelch13* resistance alleles with different amino acid mutations clearly demonstrate independent origins of artemisinin resistance. However, it is also possible that particular amino acid mutations at *kelch13* have arisen more than once: such independent alleles are expected to have different flanking SNP haplotypes. We used minimum spanning networks to examine the relationships between haplotypes surrounding different ART-R alleles to evaluate the evidence for multiple origins of particular amino acid mutations (Fig. S6). These analyses focused on the 33 SNPs immediately flanking *kelch13* in a 472kb region (1459805-1931812bp) (see Fig. S5). Three ART-R SNPs (C580Y, N458Y, and R561H) are found on divergent genotypes, suggesting 2-4 independent origins of each codon change. E252Q haplotypes are widely distributed across the minimum spanning networks. E252 haplotypes also show limited LD (Fig. 3A); for this ART-R SNP distinguishing between multiple origins and recombination is problematic.

### Estimation of the mutational target size

The mutation target size (*t*) can be directly estimated from the number of mutations observed in *kelch13* that alter parasite clearance rate. We observe 32 different non-synonymous mutations in this study, of which 26 had associated clearance rate data, and 24 showed slower clearance than WT parasites. The cumulative number observed in several published SE Asian studies (16, 18, 36–39) is 75. Note that this is a minimum estimate (*t* ≥ 75), because it includes only those sampled to date: Rarefaction methods (32), based on the rate of sampling new mutations (Fig. S7, Table S2) estimate *t* to be 106.99 (95% CI: 86.57-163.47).

### Estimation of population mutation parameter

We estimated the Θ (Pr Θ) given the observed numbers of independent ART-R alleles in population samples from 2008-14 (Fig. 4). Θ declines from 5.69 (95% CI: 3.32 – 10.61) in 2008 to 3.38 (95% CI: 1.76 – 7.34) in 2014 on the Thai-Myanmar border. Similar values are obtained from published studies of *kelch13* allele frequencies from other locations in SE Asia (Table S3). Note that these are lower bounds, because we have not accounted for resistance alleles that have arisen more than once.

**Fig. 4.**
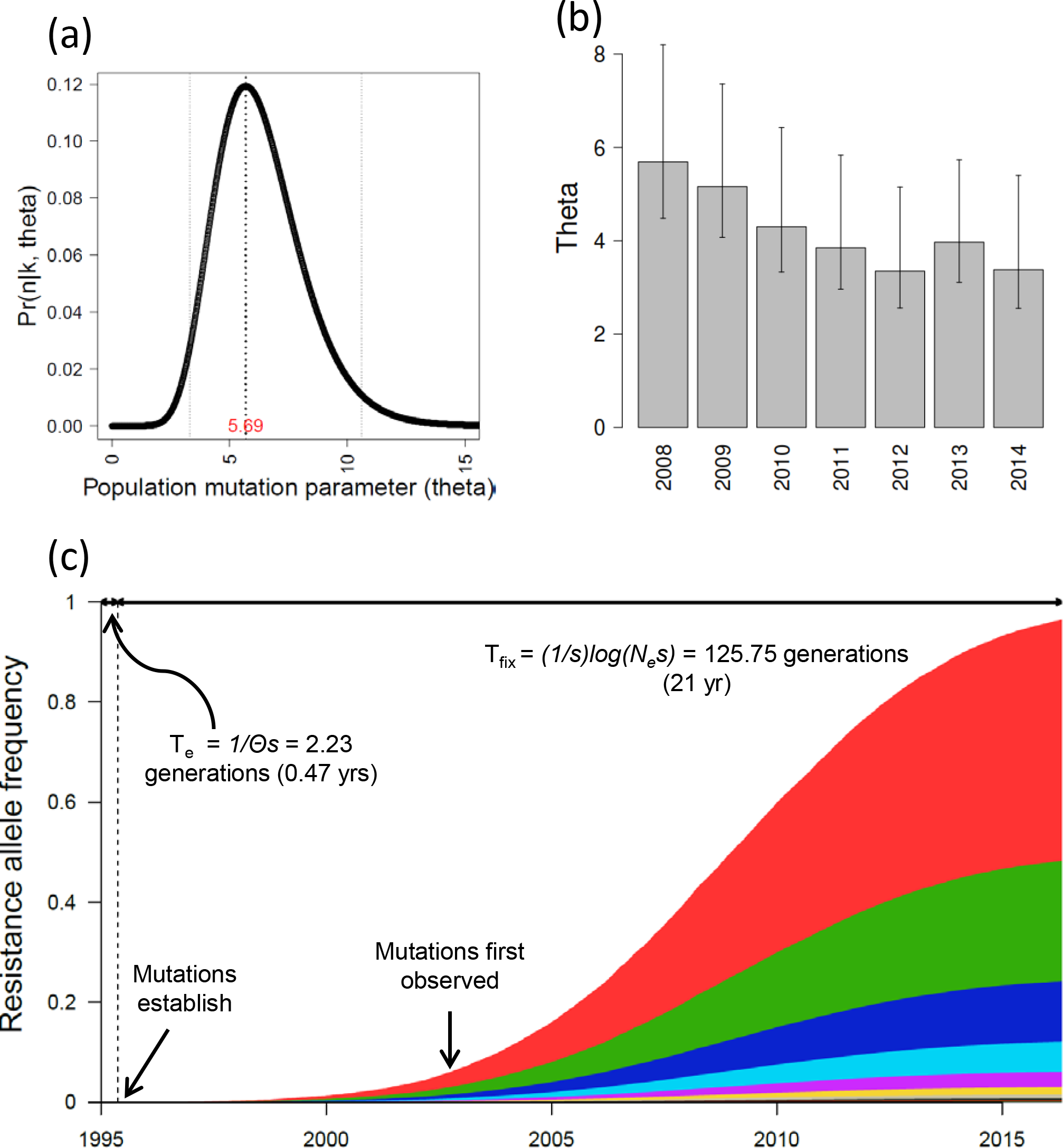
Population mutation parameter estimation. A. We estimated *Θ* (=2*N_e_μt*) from the number of different resistant alleles found in a sample of parasites from the Thai-Myanmar border (see text for details). The dotted lines show the estimate (marked in red text), and the 95% confidence errors. The data shown here is from 2008. B. *Θ* (and 95% C.I.) plotted between 2008 and 2014. *Θ* declines during this period. C. Predicted trajectory of *kelch13* alleles on the Thai-Myanmar border based on *Θ* and selection estimates (s = 0.085) from the current dataset. These predictions make the simplifying assumption that all emerging resistance alleles have equivalent fitness. ART combination therapy was assumed to be introduced in 1995, while mutations were first observed in 2003. The parameter values are those listed in Table 1, while s = 0.079. Time to establishment (*T_e_*) and fixation (T_fix)_ are estimated following Messer and Petrov (4). The dotted line shows the time when ART-R alleles are first expected to establish in the population: this is less than a year after ART introduction.

## DISCUSSION

### Mutations outside the propeller domain are associated with slow clearance

The current molecular screening approach for ART-R involves sequencing the propeller region of *kelch13* (1,725,980-1,726,940bp, amino acid positions 419-707) as this is where non-synonymous mutations associated with slow clearance rate were originally discovered in Cambodian parasite populations (40). On the Thailand-Myanmar border, the most common mutation observed prior to 2011 was E252Q. This mutation is situated 167 amino acids 5’ of the propeller region and is associated with significantly slower clearance than WT parasites, as well as significantly elevated treatment failure rates (22). Furthermore, two other mutations (D281V and R239Q) close by are also associated with slow clearance rate, arguing that molecular surveillance for resistance should be expanded outside the *kelch13* propeller domain. Furthermore, these mutations outside the propeller region may also provide valuable clues as to which sections of *kelch13* are involved in molecular interactions with other *P. falciparum* proteins. Whether these mutations are causative or linked to nearby non-coding changes associated with slow clearance is currently unclear.

Of 25 non-synonymous mutations for which accompanying clearance rate data was available, 23/25 (92%) were associated with slower clearance than WT parasites: the vast majority of mutations in this conserved gene influence phenotype. However, mutations differed significantly in associated clearance rate. E252Q shows slower clearance than WT alleles, but significantly faster clearance than the majority of propeller mutations including C580Y. Alleles also show heterogeneity in their impact on treatment failure (22), which should be directly linked to their capacity to spread. Phenotypic heterogeneity among alleles suggests that ART-R mutations in *kelch13* cause partial rather than complete loss of function. C580Y is of special interest as this allele is spreading to fixation in this population, and other studied SE Asian parasite populations (16, 40). C580Y is associated with faster clearance rate than two other mutations (R561H and N458Y). However, this difference is not significant after correction for multiple testing. The success of this allele may stem from the fact that the structural changes result from internal disruption of a disulphide bond, rather than changes in size or charge of amino acids exposed on the *kelch13* surface.

One explanation for heterogeneity in clearance rate might be involvement of other loci in the genome. For example, Miotto et al. (41) have suggested that alleles in four different genome regions may provide a permissive background for ART resistance. We found no evidence for potential involvement of genetic background on clearance rates for several common *kelch13* genotypes by comparing within and between clone variance in clearance rate. We emphasize that this does not rule out involvement of other loci in other aspects of resistance phenotype such as parasite fitness. A longitudinal genome sequencing study on a subset of the parasites examined in this paper identified several regions of the genome that show rapid changes in SNP frequency, identifying several other potential genome regions that may be selected by ART selection (42). Similarly, previous association analyses have identified other genome regions associated with ART-R other than the chr. 13 region containing *kelch13* (28, 29, 41, 43, 44).

We observed 32 non-synonymous mutations in this population. If we consider mutations observed in other SE Asian populations there are 75 non-synonymous mutations. Hence, the mutational target size is ≥75, and rarefaction analyses suggest the total number of positions in which mutations can generate a resistance phenotype is estimated at 107 (95% CI: 87-163). These include several codons in which independent mutations have occurred (P667->Q,R or T, while P527->H or L). Furthermore, examination of flanking SNPs clearly indicates independent origins of E252Q and C580Y mutations in this population. This is consistent with Takala-Harrison et al. (18), who identified independent origins of *kelch13* mutations in different SE Asian countries.

### Retrospective studies underestimate complexity of selection events

*Kelch13* heterozygosity rises until 2012 then decreases in the following two years as C580Y reaches high frequency displacing other alleles (Fig. 1E, 4B). Published datasets from Western Cambodia also support a model of diminishing variation at *kelch13* over time. As *kelch13* resistance alleles reached high frequencies in western Cambodia several years before the Thailand-Myanmar border, they provide a snapshot of what we expect will occur on the Thai-Myanmar border in future years. For example, *kelch13* heterozygosity has also diminished in parasites sampled from Pursat, Battambang and Pailin in western Cambodia between 2001-12 (40).

Diminishing complexity of selective events at dihydrofolate reductase (*dhfr*) is also observed during evolution of pyrimethamine resistance in both Africa and Asia. For example, on the Thai-Myanmar border, there were several independent origins of S108N mutation, which is the first step of resistance evolution (10). However, high level resistance involving N51I, C59R and I164L was only associated with a single chr. 4 haplotype. Similarly, in Africa, there were multiple origins of parasites containing a single mutation (S108N), three separate origins of parasites carrying double mutant (N51I/S108N or C59R/S108N) (45), but these have been rapidly replaced by triple mutant parasites originating in Asia (11).

Both *kelch13* and *dhfr* datasets support a model in which selection events are soft initially, but rapidly become harder, as particular alleles dominate. Continuing surveillance of the parasite population on the Thai-Myanmar border will determine whether this trend continues and eventually leaves a single resistance *kelch13* mutation and a hard signature. Most theoretical work assumes that beneficial alleles selected during soft selective sweeps are functionally equivalent, and therefore whether particular beneficial mutations fix is dependent on genetic drift rather than selection (3). Demographic alone can result in loss of adaptive alleles (46). However, in the malaria examples, differences in the selection coefficients of resistance alleles are most likely responsible for the hardening of the sweep. *Kelch13* alleles are not functionally equivalent and differ in both clearance and ability to survive treatment (22). That independent C580Y alleles have spread to high frequency in different countries in SE Asia also suggests that this allele has higher fitness than other ART-R alleles. More generally, we suggest that retrospective studies of sequence variation in extant populations are likely to underestimate the frequency and complexity of soft selective events.

An interesting feature of these data is that multiple alleles showing moderate fitness benefit initially spread in this population. For example the E252Q allele is associated with a modest change in clearance rate, and a modest impact on surviving ART-treatment relative to other ART-alleles. C580Y out competes other alleles after 2011. These data are consistent with both theoretical and empirical work on adaptation demonstrating that the distribution of fitness effects of beneficial mutations is exponential: while multiple mutations can generate phenotypes with modest increase in fitness, only a small subset of mutations show high fitness (47, 48). The time taken for such high fitness alleles to arise and establish is expected to be greater than for alleles of moderate fitness. The rarity of high fitness ART-R alleles may explain why the C580Y allele appeared after other alleles, but subsequently swept through the population on the Thai-Myanmar border.

### Population mutation parameter and *N_e_* estimates

*N_e_* is a central measure in population genetic analysis that determines the level of genetic drift or inbreeding in idealized populations (49), is key to understanding the rate at which adaptive mutations are introduced into populations (3, 4), but is difficult to measure directly. In this case we are interested in understanding short term *N_e_* as this is the period in which ART selection is operating (from 1995 onwards). *N_e_* can be inferred for natural populations by measuring variation in allele frequencies over time (i.e. by quantifying genetic drift). This approach has been used in the parasite population studied on the Thai-Myanmar border, providing average *N_e_* estimates of 1965 (14) and in Senegal where *N_e_* estimates were <100. We can also infer the likely value of Θ from the numbers of independent adaptive alleles observed in our population sample. This approach to estimation of *N_e_* from *Θ* follows work on *Drosophila* (6) and HIV (5) and utilizes the fact that *Θ*=2*N_e_μ* (and therefore *N_e_*=*Θ*/2*μ*). Θ ranges from 5.9 to 3.9 between 2008-14 on the Thai-Myanmar border and comparable calculations from other SE Asian locations provide remarkably similar estimates (Table S3). Using estimates of minimum and maximum mutation rate estimates (5.10 × 10^−8^ to 1.15 ×10^−7^ per nucleotide per generation) and target size (87– 163 bp), we determined short term *N_e_* to be from 88,017 – 1,195,628 on the Thai-Myanmar border (Table 1). These estimates of *N_e_* are 52-705 times greater than previous estimates of *Ne* from the same parasite population based on fluctuation in allele frequencies at 96 SNPs between 2001-10 (14). These values are also 9-120 times greater than long-term *N_e_* estimates (~10,000) in SE Asia based on nucleotide diversity in mtDNA (50). One caveat is that we may have underestimated the mutational target size (*t*). To provide a generous upper bound on the target size we can assume that mutations in all 1771 non-synonymous sites within *kelch13* can generate resistance alleles. Even with this obvious overestimate of *t*, *N_e_* estimates (*N_e_* = 15,479) are still 9-fold greater than variance *N_e_* values from the same population.

**Table 1.** Summary of parameters used for estimation of population mutation parameter (*Θ*) and *N_e_*

| Parameter | Estimate | Lower bound | Upper bound | Notes | Citation |
| --- | --- | --- | --- | --- | --- |
| **Mutation rate/base/48 hr asexual cycle** | 3.39E-09 | 1.7E-09 | 3.82E-09 | Average values from 5 mutation accumulation experiments: upper and lower bounds are highest and lowest values recorded | (30, 31) |
| **Mutation rate/base/generation ($\mu$)** | 1.02E-07 | 5.10E-08 | 1.15E-07 | Assumes 2 months generation time (mosquito to mosquito). | (30, 31) |
| **Mutation target size (bp) ($t$)**<br>The number of base changes in *kelch13* that can result in Artemisinin resistance | 106.99 | 86.57 | 163.47 | Estimates derived by rarefaction (Table S2, Fig. S7) using published datasets and this study | (16, 18, 36-39) |
| **Population mutation parameter ($\Theta$)** | 5.69 | 3.32 | 10.61 | Estimated using eqn. 12 in Pennings and Hermisson (2005) from observed number of *kelch13* alleles ($k$) in samples of size $n$ from the Thai-Myanmar border (Fig. 4). Values from 2008 shown | This study |
| **$N_e$ (estimated from $\Theta$)** | 2.61E+05 | 8.86E+04 | 1.20E+06 | Estimated from the parameters listed above: $N_e = \Theta/2\mu t$ | This study |

Two recent papers have challenged the application of classical population genetics models to malaria parasites and other pathogens (51, 52). Classical population genetics models make Wright-Fisher assumptions that populations are constant in size, have non-overlapping generations, and that each new generation is sampled from gametes produced by the previous generation. In contrast, the malaria parasite lifecycle consists of very large populations (≤10^11^) of asexually dividing blood stage forms (“gametes”) within humans, from which small numbers of gametes (≤10) are sampled by mosquitoes to found the following generation. Simulation studies demonstrate that under the malaria lifecycle, the role of genetic drift is amplified relative to the Wright-Fisher model because small numbers of gametes are sampled from infected people, while the role of selection is amplified because there are very large numbers of parasites within patients (52). Hence, drift-based measures of *N_e_*, using fluctuation in allele frequencies, reflect most closely the numbers of parasites passing through the mosquito bottleneck, and provide a gross underestimate of the parasite population relevant for adaptation. We suggest that this decoupling of drift and selection in the malaria lifecycle, may explain the discrepancy between *N_e_* estimates, and that the *N_e_* relevant for adaptation can best be estimated from *Θ*. Similar lifecycles are found in multiple pathogens and parasites: more thorough evaluation of the potential bias in applying classical population genetic tools to these organisms is urgently needed. Previous analyses have suggested that the availability of mutations limits drug resistance evolution in *Plasmodium* (53). The large *N_e_* values inferred here suggest we have underestimated the capacity for adaptive evolution in *Plasmodium*.

### A spectrum of selective events in Plasmodium

Selective events associated with 5 additional loci have been extensively studied in SE Asian parasite populations and reveal a spectrum of sweeps ranging from hard to soft. We use estimates of *Θ* to better understand this spectrum. Both the chloroquine resistance transporter (*crt*) and *dhfr* provide examples of hard selective sweeps in SE Asia. A single allele (K76T) at *crt* spread through SE Asian malaria populations, purging genetic variation from genomic regions flanking *crt*, and this allele subsequently also spread across Africa (12). In the case of *dhfr*, 2-4 mutations conferring resistance have been observed on the Thai-Myanmar border. However, all of these mutations share the same flanking regions, consistent with a series of hard selective sweeps in which mutations arose successively on the same sweeping haplotype (10). Resistance at both *crt* and *dhfr* involves specific single SNP mutations, so the target size (t) is 1. As a consequence Θ will be reduced ~100-fold relative to *kelch13* to ~0.05. In this range of Θ values, multiple coexisting alleles are expected to be rare. Hence, the observation of hard sweeps at *dhfr* and *crt* is broadly consistent with and dependent on target size. Resistance alleles at multidrug transporter (*mdr1) and GTP cyclohydrolase I (gch1*) and dihydropteroate synthase (*dhps*) loci have multiple independent origins. Both *mdr1* and *gch1* involve multiple independent origins of resistance-associated copy number variants (CNVs). There are between 5-15 independent origins of resistance-associated CNVs containing *mdr1* (54), and at least three independent origins of CNVs containing *gch1* (55). In this case, the elevated mutation rate of CNV compared with SNPs most likely explains the multiple co-existing alleles. Finally, studies of *dhps* evolution, which involves five specific amino acid positions, strongly suggest several origins, because microsatellite loci flanking this locus show minimal reduction in variation relative to wild-type alleles and several different haplotypes are associated with resistant *dhps* alleles (56–58).

### Implications for resistance evolution and management

Our re-estimation of *N_e_* for malaria in SE Asia has important implications for resistance evolution and management. *ART* combination therapy was introduced on the Thai-Myanmar border in 1995, resistance was first suspected in 2008, and resistance alleles have now almost reached fixation. When did resistance alleles first become established in this parasite population? The time to establishment (*T_e_*) – when resistance alleles first escape elimination by genetic drift – is: *T_e_* = 1/*πs* (4). Hence, *T_e_* is 2.23 generations or 4.66 months (assuming 6 generations/yr) for this parasite population, suggesting that resistance alleles likely became established on the Thai-Myanmar border in the mid-nineties within a year of *ART* deployment. This is 8 years before resistance alleles were first observed in this longitudinal dataset (Fig. 1 and 4), and 13 years before resistance was suspected in SE Asia (17). The average number of ART-R mutations arising between 1995 and fixation is 2*θN_e_log(α)/α*, where *α* = 2*N_e_s* (3). Hence, with our estimated parameters (Table 1) ART-R mutations will have arisen 716 times in the course of this selective event.

Artemisinin is used to treat patients in combination with other drugs such as lumefantrine, mefloquine and piperaquine. This is expected to constrain resistance evolution, because the probabilities of a parasite gaining mutations against two different drugs is the product of the rate at which mutation generates resistance alleles conferring resistance against each individual drug. This policy appears to have worked poorly in restraining resistance evolution given that ART resistance has evolved multiple times, and ART-R mutations are estimated to have established in treated population <1 year following introduction of ART combinations. The use of partner drugs (e.g. mefloquine and lumefrantine) to which resistance has already emerged is one contributor to the failure of ACTs. Our results suggest that antimalarial combinations will need to contain more than two fully effective drugs to effectively constrain resistance evolution. This conclusion is very similar to that reached by HIV researchers (59). Resistance breakthroughs occurred rapidly within patients treated with early HIV drug cocktails with multiple origins of resistance occurring within single infections. As HIV drug combinations increased in complexity, drug breakthroughs have became increasingly rare and drug resistance associated sweeps significantly harder (60). In South East Asia triple combination containing artesunate and mefloquine + piperaquine, or lumefantrine + amodiaquine are now under clinical trials. However, use of triple or quadruple combinations is severely limited by the availability of possible drugs.

The evolution of ART-R demonstrates the importance of mutational target size for resistance evolution. We suggest that target size can readily be incorporated into preliminary evaluations of promising new antimalarial compounds, prior to deployment. Laboratory selection of cultured parasites using new antimalarials is now routinely carried out to evaluate the potential for resistance evolution (61). Such studies can be used both to identify genes that underlie resistance, but also to determine how many nucleotides can generate resistance phenotype. An exciting approach to target size evaluation might be to use CRISPR/Cas9 approaches with degenerate homology regions to generate a population of mutations across a target gene, and then evaluate what fraction of these mutations responds to drug selection. In essence, this approach increases mutation rate in the target gene to allow thorough evaluation of mutational target size.

## Acknowledgements

We thank National Institutes for Health grant R37 AI048071 (TJCA) and the Bill and Melinda Gates Foundation for funding. Comments from Pleuni Pennings, Philipp Messer, and Dan Neafsey improved the manuscript. This work was conducted in facilities constructed with support from Research Facilities Improvement Program grant C06 RR013556 from the National Center for Research Resources. SMRU is part of the Mahidol Oxford University Research Unit supported by the Wellcome Trust of Great Britain.

## References

1. Schrider DR, Mendes FK, Hahn MW, Kern AD. Soft shoulders ahead: spurious signatures of soft and partial selective sweeps result from linked hard sweeps. Genetics. 2015 May;200(1):267–84.

2. Jensen JD. On the unfounded enthusiasm for soft selective sweeps. Nat Commun. 2014 Oct 27;5:5281.

3. Pennings PS, Hermisson J. Soft sweeps II--molecular population genetics of adaptation from recurrent mutation or migration. Mol Biol Evol. 2006 May;23(5):1076–84.

4. Messer PW, Petrov DA. Population genomics of rapid adaptation by soft selective sweeps. Trends Ecol Evol. 2013 Nov;28(11):659–69.

5. Pennings PS, Kryazhimskiy S, Wakeley J. Loss and recovery of genetic diversity in adapting populations of HIV. PLoS Genet. 2014 Jan;10(1):e1004000.

6. Karasov T, Messer PW, Petrov DA. Evidence that adaptation in Drosophila is not limited by mutation at single sites. PLoS Genet. 2010 Jun 17;6(6):e1000924.

7. Barroso-Batista J, Sousa A, Lourenco M, Bergman ML, Sobral D, Demengeot J, et al. The first steps of adaptation of Escherichia coli to the gut are dominated by soft sweeps. PLoS Genet. 2014 Mar 6;10(3):e1004182.

8. Weetman D, Mitchell SN, Wilding CS, Birks DP, Yawson AE, Essandoh J, et al. Contemporary evolution of resistance at the major insecticide target site gene Ace-1 by mutation and copy number variation in the malaria mosquito Anopheles gambiae. Mol Ecol. 2015 Jun;24(11):2656–72.

9. Anderson T, Nkhoma S, Ecker A, Fidock D. How can we identify parasite genes that underlie antimalarial drug resistance? Pharmacogenomics. 2011 Jan;12(1):59–85.

10. Nair S, Williams JT, Brockman A, Paiphun L, Mayxay M, Newton PN, et al. A selective sweep driven by pyrimethamine treatment in southeast asian malaria parasites. Mol Biol Evol. 2003 Sep;20(9):1526–36.

11. Roper C, Pearce R, Nair S, Sharp B, Nosten F, Anderson T. Intercontinental spread of pyrimethamine-resistant malaria. Science. 2004 Aug 20;305(5687):1124.

12. Wootton JC, Feng X, Ferdig MT, Cooper RA, Mu J, Baruch DI, et al. Genetic diversity and chloroquine selective sweeps in Plasmodium falciparum. Nature. 2002 Jul 18;418(6895):320–3.

13. Chang HH, Park DJ, Galinsky KJ, Schaffner SF, Ndiaye D, Ndir O, et al. Genomic sequencing of Plasmodium falciparum malaria parasites from Senegal reveals the demographic history of the population. Mol Biol Evol. 2012 Nov;29(11):3427–39.

14. Nkhoma SC, Nair S, Al-Saai S, Ashley E, McGready R, Phyo AP, et al. Population genetic correlates of declining transmission in a human pathogen. Mol Ecol. 2013 Jan;22(2):273–85.

15. Sibley CH. Malaria historians: analyze your old slides! Trends Parasitol. 2006 Jun;22(6):243–4.

16. Ashley EA, Dhorda M, Fairhurst RM, Amaratunga C, Lim P, Suon S, et al. Spread of artemisinin resistance in Plasmodium falciparum malaria. N Engl J Med. 2014 Jul 31;371(5):411–23.

17. Dondorp AM, Nosten F, Yi P, Das D, Phyo AP, Tarning J, et al. Artemisinin resistance in Plasmodium falciparum malaria. N Engl J Med. 2009 Jul 30;361(5):455–67.

18. Takala-Harrison S, Jacob CG, Arze C, Cummings MP, Silva JC, Dondorp AM, et al. Independent emergence of artemisinin resistance mutations among Plasmodium falciparum in Southeast Asia. J Infect Dis. 2015 Mar 1;211(5):670–9.

19. Straimer J, Gnadig NF, Witkowski B, Amaratunga C, Duru V, Ramadani AP, et al. Drug resistance. K13-propeller mutations confer artemisinin resistance in Plasmodium falciparum clinical isolates. Science. 2015 Jan 23;347(6220):428–31.

20. Spring MD, Lin JT, Manning JE, Vanachayangkul P, Somethy S, Bun R, et al. Dihydroartemisinin-piperaquine failure associated with a triple mutant including kelch13 C580Y in Cambodia: an observational cohort study. Lancet Infect Dis. 2015 Jun;15(6):683–91.

21. Amaratunga C, Lim P, Suon S, Sreng S, Mao S, Sopha C, et al. Dihydroartemisinin-piperaquine resistance in Plasmodium falciparum malaria in Cambodia: a multisite prospective cohort study. Lancet Infect Dis. 2016 Jan 7.

22. Phyo AP, Ashley EA, Anderson TJ, Bozdech Z, Carrara VI, Sriprawat K, et al. Declining efficacy of artemisinin combination therapy against P. falciparum malaria on the Thai-Myanmar border (2003-2013): the role of parasite population genetic factors. Clin.Infect.Dis. In Press.

23. Venkatesan M, Amaratunga C, Campino S, Auburn S, Koch O, Lim P, et al. Using CF11 cellulose columns to inexpensively and effectively remove human DNA from Plasmodium falciparum-infected whole blood samples. Malar J. 2012 Feb 10;11:41,2875-11-41.

24. Phyo AP, Nkhoma S, Stepniewska K, Ashley EA, Nair S, McGready R, et al. Emergence of artemisinin-resistant malaria on the western border of Thailand: a longitudinal study. Lancet. 2012 May 26;379(9830):1960–6.

25. Carrara VI, Lwin KM, Phyo AP, Ashley E, Wiladphaingern J, Sriprawat K, et al. Malaria burden and artemisinin resistance in the mobile and migrant population on the Thai-Myanmar border, 1999-2011: an observational study. PLoS Med. 2013;10(3):e1001398.

26. Dondorp AM, Yeung S, White L, Nguon C, Day NP, Socheat D, et al. Artemisinin resistance: current status and scenarios for containment. Nat Rev Microbiol. 2010 Apr;8(4):272–80.

27. Flegg JA, Guerin PJ, White NJ, Stepniewska K. Standardizing the measurement of parasite clearance in falciparum malaria: the parasite clearance estimator. Malar J. 2011 Nov 10;10:339,2875-10-339.

28. Cheeseman IH, Miller BA, Nair S, Nkhoma S, Tan A, Tan JC, et al. A major genome region underlying artemisinin resistance in malaria. Science. 2012 Apr 6;336(6077):79–82.

29. Cheeseman IH, McDew-White M, Phyo AP, Sriprawat K, Nosten F, Anderson TJ. Pooled sequencing and rare variant association tests for identifying the determinants of emerging drug resistance in malaria parasites. Mol Biol Evol. 2015 Apr;32(4):1080–90.

30. Claessens A, Hamilton WL, Kekre M, Otto TD, Faizullabhoy A, Rayner JC, et al. Generation of antigenic diversity in Plasmodium falciparum by structured rearrangement of Var genes during mitosis. PLoS Genet. 2014 Dec 18;10(12):e1004812.

31. Bopp SE, Manary MJ, Bright AT, Johnston GL, Dharia NV, Luna FL, et al. Mitotic evolution of Plasmodium falciparum shows a stable core genome but recombination in antigen families. PLoS Genet. 2013;9(2):e1003293.

32. Colwell RK. EstimateS: statistical estimation of species richness and shared species from samples. 2013;Version 9.

33. Gautier M, Vitalis R. rehh: an R package to detect footprints of selection in genome-wide SNP data from haplotype structure. Bioinformatics. 2012 Apr 15;28(8):1176–7.

34. Neafsey DE, Schaffner SF, Volkman SK, Park D, Montgomery P, Milner DA,Jr, et al. Genome-wide SNP genotyping highlights the role of natural selection in Plasmodium falciparum population divergence. Genome Biol. 2008;9(12):R171,2008-9-12-r171. Epub 2008 Dec 15.

35. Kamvar ZN, Tabima JF, Grunwald NJ. Poppr: an R package for genetic analysis of populations with clonal, partially clonal, and/or sexual reproduction. PeerJ. 2014 Mar 4;2:e281.

36. Nyunt MH, Hlaing T, Oo HW, Tin-Oo LL, Phway HP, Wang B, et al. Molecular assessment of artemisinin resistance markers, polymorphisms in the k13 propeller, and a multidrug-resistance gene in the eastern and western border areas of Myanmar. Clin Infect Dis. 2015 Apr 15;60(8):1208–15.

37. Wang Z, Wang Y, Cabrera M, Zhang Y, Gupta B, Wu Y, et al. Artemisinin Resistance at the China-Myanmar Border and Association with Mutations in the K13 Propeller Gene. Antimicrob Agents Chemother. 2015 Nov;59(11):6952–9.

38. Tun KM, Imwong M, Lwin KM, Win AA, Hlaing TM, Hlaing T, et al. Spread of artemisinin-resistant Plasmodium falciparum in Myanmar: a cross-sectional survey of the K13 molecular marker. Lancet Infect Dis. 2015 Apr;15(4):415–21.

39. Huang F, Takala-Harrison S, Jacob CG, Liu H, Sun X, Yang H, et al. A Single Mutation in K13 Predominates in Southern China and Is Associated With Delayed Clearance of Plasmodium falciparum Following Artemisinin Treatment. J Infect Dis. 2015 Nov 15;212(10):1629–35.

40. Ariey F, Witkowski B, Amaratunga C, Beghain J, Langlois AC, Khim N, et al. A molecular marker of artemisinin-resistant Plasmodium falciparum malaria. Nature. 2014 Jan 2;505(7481):50–5.

41. Miotto O, Amato R, Ashley EA, MacInnis B, Almagro-Garcia J, Amaratunga C, et al. Genetic architecture of artemisinin-resistant Plasmodium falciparum. Nat Genet. 2015 Mar;47(3):226–34.

42. Cerqueira GC, Cheeseman IH, Nair S, McDew-White M, Melnikov A, Nosten F, et al. Retrospective longitudinal genomic surveillance of Plasmodium falciparum malaria parasites documents the emergence of artemisinin resistance in Thailand. October 25-29; Annual Meeting of the American Society of Tropical Medicine and Hygiene, Philadelphia. ; 2015.

43. Takala-Harrison S, Clark TG, Jacob CG, Cummings MP, Miotto O, Dondorp AM, et al. Genetic loci associated with delayed clearance of Plasmodium falciparum following artemisinin treatment in Southeast Asia. Proc Natl Acad Sci U S A. 2013 Jan 2;110(1):240–5.

44. Miotto O, Almagro-Garcia J, Manske M, Macinnis B, Campino S, Rockett KA, et al. Multiple populations of artemisinin-resistant Plasmodium falciparum in Cambodia. Nat Genet. 2013 Jun;45(6):648–55.

45. Roper C, Pearce R, Bredenkamp B, Gumede J, Drakeley C, Mosha F, et al. Antifolate antimalarial resistance in southeast Africa: a population-based analysis. Lancet. 2003 Apr 5;361(9364):1174–81.

46. Wilson BA, Petrov DA, Messer PW. Soft selective sweeps in complex demographic scenarios. Genetics. 2014 Oct;198(2):669–84.

47. Kassen R, Bataillon T. Distribution of fitness effects among beneficial mutations before selection in experimental populations of bacteria. Nat Genet. 2006 Apr;38(4):484–8.

48. Orr HA. The distribution of fitness effects among beneficial mutations. Genetics. 2003 Apr;163(4):1519–26.

49. Charlesworth B. Fundamental concepts in genetics: effective population size and patterns of molecular evolution and variation. Nat Rev Genet. 2009 Mar;10(3):195–205.

50. Joy DA, Feng X, Mu J, Furuya T, Chotivanich K, Krettli AU, et al. Early origin and recent expansion of Plasmodium falciparum. Science. 2003 Apr 11;300(5617):318–21.

51. Chang HH, Hartl DL. Recurrent bottlenecks in the malaria life cycle obscure signals of positive selection. Parasitology. 2015 Feb;142 Suppl 1:S98–S107.

52. Chang HH, Moss EL, Park DJ, Ndiaye D, Mboup S, Volkman SK, et al. Malaria life cycle intensifies both natural selection and random genetic drift. Proc Natl Acad Sci U S A1. 2013 Dec 10;110(50):20129–34.

53. Anderson TJ, Roper C. The origins and spread of antimalarial drug resistance: lessons for policy makers. Acta Trop. 2005 Jun;94(3):269–80.

54. Nair S, Nash D, Sudimack D, Jaidee A, Barends M, Uhlemann AC, et al. Recurrent gene amplification and soft selective sweeps during evolution of multidrug resistance in malaria parasites. Mol Biol Evol. 2007 Feb;24(2):562–73.

55. Nair S, Miller B, Barends M, Jaidee A, Patel J, Mayxay M, et al. Adaptive copy number evolution in malaria parasites. PLoS Genet. 2008 Oct;4(10):e1000243.

56. Vinayak S, Alam MT, Mixson-Hayden T, McCollum AM, Sem R, Shah NK, et al. Origin and evolution of sulfadoxine resistant Plasmodium falciparum. PLoS Pathog. 2010 Mar 26;6(3):e1000830.

57. Pearce RJ, Pota H, Evehe MS, Ba e, Mombo-Ngoma G, Malisa AL, et al. Multiple origins and regional dispersal of resistant dhps in African Plasmodium falciparum malaria. PLoS Med. 2009 Apr 14;6(4):e1000055.

58. Naidoo I, Roper C. Following the path of most resistance: dhps K540E dispersal in African Plasmodium falciparum. Trends Parasitol. 2010 Sep;26(9):447–56.

59. Pennings PS. HIV Drug Resistance: Problems and Perspectives. Infect Dis Rep..2013 Jun 6;5(Suppl 1):e5.

60. Feder AF, Rhee SY, Holmes SP, Shafer RW, Petrov DA, Pennings PS. More effective drugs lead to harder selective sweeps in the evolution of drug resistance in HIV-1. Elife. 2016 Feb 15;5:10.7554/eLife.10670.

61. Flannery EL, Fidock DA, Winzeler EA. Using genetic methods to define the targets of compounds with antimalarial activity. J Med Chem. 2013 Oct 24;56(20):7761–71.

